# Towards Automated Acoustic Monitoring of Threatened Birds in Tropical Forest Ecosystem Restoration

**DOI:** 10.64898/2026.09.06.749699

**Authors:** Heather X. Fortune, Rhys Preston-Allen, Xiuhan Zhang, Celso Henrique de Freitas Parruco, Cristina Banks-Leite

## Abstract

Forest restoration has become a widespread response to deforestation and biodiversity loss, particularly across the tropics. Monitoring restoration is essential for tracking progress and requires accurate knowledge of baseline communities, with threatened species often used as recovery indicators. Passive Acoustic Monitoring (PAM) combined with AI classifiers is enabling this at scale. However, such tools remain immature and may underperform for rare, threatened species underrepresented in training data. Here, we demonstrate the potential of PAM assisted by the AI classifier BirdNET+ V3.0, a January 2026 beta release, to support monitoring of threatened tropical birds. By validating 995 detections from 8,940 hours of audio across a reforestation landscape in Pará in the Brazilian Amazon, we found that BirdNET+ confirmed the presence of ∼43% of the IUCN Red List threatened species possible in the region, five of which were the first confirmed records for the study site. We also found that this method outperformed traditional survey methods. However, BirdNET+ identified species with mixed reliability, and we could also not determine whether restoration has had any impact on them. Our results demonstrate that PAM with BirdNET+ is a viable, practical method for detecting threatened birds in tropical forest restoration landscapes, but that conventional validation approaches are not appropriate for such species, for which extensive validation effort is an essential cost. Under circumstances where this is not possible, we demonstrate validating the single highest-confidence detection per occasion as a practical alternative.

## Introduction

Deforestation remains a key driver of the ongoing climate and biodiversity crises (Betts et al., 2017). Forest restoration has emerged as a global response, with millions of hectares to be restored worldwide within the next decade - a result of efforts like the Bonn Challenge (International Union for Conservation of Nature, 2020). The potential of such initiatives extends beyond conservation to the active recovery of biodiversity and ecosystem function (Derhé et al., 2016; Kemppinen et al., 2020). Tropical rainforests have received special attention owing to elevated deforestation rates, alongside their significant levels of biodiversity, endemism, and carbon storage potential (Lapola et al., 2023). With more than a tenth of the world’s terrestrial biodiversity, the Amazon rainforest has been a key focus (Science Panel for the Amazon, 2021). In the Brazilian Amazon alone, over four hundred restoration projects have been established as of 2017 (da Cruz et al., 2021).

However, prior to any measurable biodiversity recovery, the establishment phase of a forest restoration site can impose a substantial disturbance event, requiring the construction of access roads, transport of materials, site preparation, alongside the planting operation itself (Axelsson et al., 2024). Anthropogenic disturbance from sources such as these has been shown to disrupt faunal communities, decreasing species richness and abundance, or causing displacement, with cascading consequences for the rest of the ecosystem (Francis, Ortega & Cruz, 2009; Sordello et al., 2020). This is particularly relevant for threatened species, whose small population sizes and restricted ranges elevate their extinction risk (Owens & Bennett, 2000; Sodhi et al., 2008). Therefore, monitoring the impacts of restoration on local biodiversity is imperative to recognise and mitigate any potential negative effects.

Beyond identifying biodiversity impacts, monitoring is also essential for tracking project progress and evaluating success, which requires accurate knowledge of baseline communities (Ruiz-Jaen & Mitchell Aide, 2005). This is important not just from a conservation perspective, but also for carbon credit schemes, where accurate baselines underpin legitimacy. In response, instruments like the Species Threat Abatement and Restoration (STAR) Framework and the Isometric Standard have emerged to standardise progress measurement, ensure credibility, and inform management decisions (Isometric, 2023; Mair et al., 2021). These typically focus on species of conservation concern, whose rarity and declining status give them the greatest potential for conservation gains (Joseph, Maloney & Possingham, 2009). Such species are also often endemic or hold cultural value, serving as surrogates for wider ecosystem conservation (Caro, 2010). They further serve as indicators of ecosystem intactness, and birds in particular - with critical ecosystem functions and a wide range of niches - are established indicator species (Fraixedas et al., 2020). However, fulfilling these purposes requires accurate data on their presence and population trends.

Restoration projects generally take a matter of decades, thus monitoring these species requires scalability across extensive spatial and temporal scales. This is costly and logistically challenging with traditional methods like point counts, which require experts to identify species, and are time- and access-constrained (Shonfield & Bayne, 2017). Passive Acoustic Monitoring (PAM) provides an alternative, where autonomous recording units (ARUs) are deployed across a landscape at fixed points and can record unattended for weeks, capturing a wide range of taxa which primarily communicate by sound (Browning et al., 2017; Sugai et al., 2019). Birds, which are wide-ranging and have species-specific vocalisations, are particularly well-suited to PAM (Inoue et al., 2025). This method has grown in popularity as it enables non-invasive monitoring, flexible and extended recording periods compared with human observers, and retrospective review of recordings (Knight et al., 2017; Zwerts et al., 2021). These factors increase the likelihood of capturing rare occurrences, making PAM effective for species of conservation concern, which are characterised by low detection counts (Manzano-Rubio et al., 2022; Sidie-Slettedahl et al., 2015).

The main challenge of PAM is that it generates vast quantities of audio data that must then be analysed to detect and identify species. As such, automated Artificial Intelligence (AI) tools, primarily Deep Learning (DL) models, have become widely adopted for this task, although they are in their infancy (Márquez-Rodríguez et al., 2025; Stowell et al., 2016). For birds, BirdNET is currently the field standard: a convolutional neural network (CNN) model developed by the Cornell Institute of Ornithology, which serves as an off-the-shelf automated AI classifier that identifies species without the need for human expertise (Kahl et al., 2021). The current stable release, V2.4, can recognise over 6000 species worldwide (Kahl et al., 2025). This open-source tool thus allows even beginners and citizen scientists to efficiently process many hours of audio data and supports repeatable monitoring free from observer bias (Usman, Alce & Galido, 2026).

BirdNET has demonstrated strong performance for common, temperate species, which is unsurprising since its main training datasets, Xeno-canto and Macaulay Library, are biased towards European and North American birds (Stowell, 2022). Consequently, it struggles to detect or correctly identify tropical species, which remain underrepresented in these databases (Sethi et al., 2024). This is compounded by the “long-tailed” nature of species distributions, meaning far fewer recordings - and thus training data - exist for rare species, which BirdNET also performs poorly for (Márquez-Rodríguez et al., 2025; Pérez-Granados et al., 2026). Given the higher levels of species richness, this effect is likely even more pronounced in the tropics (Ulrich et al., 2020).

BirdNET+ V3.0 is a beta model with an improved architecture and broader species coverage, released in January 2026 (Lasseck et al. 2026). According to the developers, it recognises over 11,000 sound classes (including a range of taxa and non-biological sounds). Of the 9,834 bird species it covers, 1853 are found in Brazil, out of the country’s 1,916 total (Lasseck et al., 2026; Lepage, 2026).

However, as a developer preview, its performance has yet to be evaluated independently in real-world settings, and there remain open questions about whether its stated improvements extend to rare species or tropical regions underrepresented in BirdNET’s training data. This is a major concern, given that so many restoration projects are concentrated in the tropics, where the threatened species being monitored are typically rare.

To establish whether PAM combined with AI classification can support the monitoring of threatened tropical birds, I conducted an independent evaluation of BirdNET+ V3.0 within a regenerating forest landscape in the Brazilian Amazon. The project is in its early stages, where reforestation has been ongoing for only three years.

The following research questions were investigated:

- RQ1: Is BirdNET+ capable of detecting threatened species in the Brazilian Amazon?
- RQ2: How does the performance of PAM using BirdNET+ compare against traditional monitoring methods in detecting threatened species?
- RQ3: Contingent on the detection capability of BirdNET+ (RQ1) and its performance relative to traditional methods (RQ2), does the community-level presence of threatened species change between baseline (T0) and after one year of restoration (T1)?

We used data from a reforestation study to test this, hypothesising a decrease in presence due to increased disturbance during the establishment phase.

## Methods

### Study Area & Design

The study site was a reforestation landscape across Pará state in northern Brazil in the Amazon (Figure 1). It consisted of several former cattle ranches undergoing restoration (n=5), as well as old-growth forest control sites (n=3) which reflect the target post-restoration state. Both site types are referred to collectively as farms throughout to avoid ambiguity with sampling points. This study covered a total of 145 fixed sampling points across these five restoration (n=89) and three control (n=56) farms, covering approximately 34,000 hectares in total. Sampling points were located a minimum of 500 metres apart – matching the effective detection radius of acoustic recorders – to limit the risk of spatial autocorrelation between neighbouring points (Browning et al., 2017; Lawson et al., 2023).

**Figure 1.**
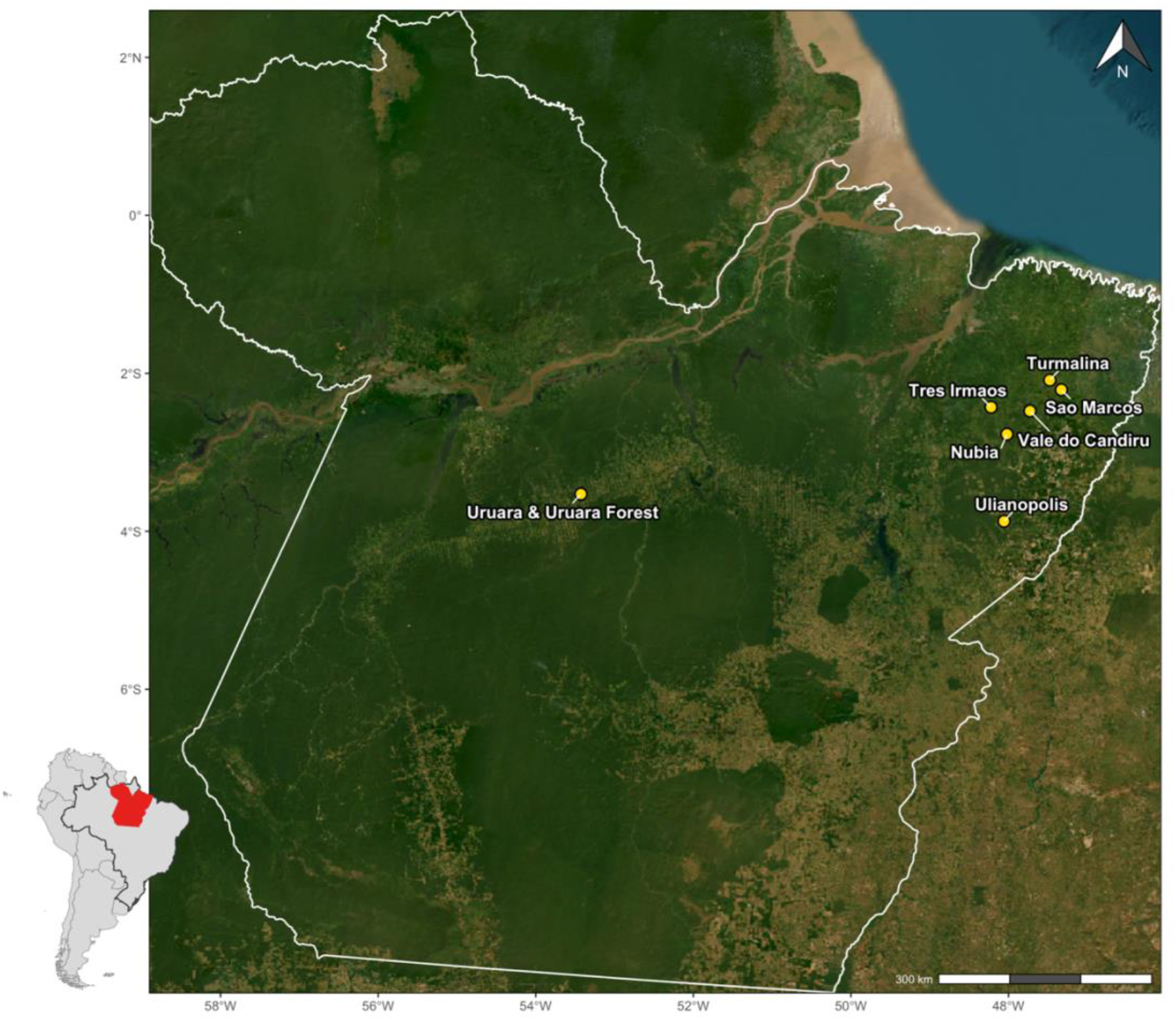
Spatial distribution of the eight study farms (restoration and forest control) across Pará state, northeastern Brazil. Farms are indicated by the yellow markers at the approximate centroid of each (individual farm boundaries are not shown at this scale). Uruará and Uruará Forest sampling points correspond to the same landholding therefore are shown as a single marker. The inset map shows the location of Pará (red) within Brazil and the wider South American continent. Generated in R (version 4.5.2; R Core Team, 2025) using the sf (Pebesma, 2018), maptiles (Giraud, 2026), and rnaturalearth (Massicotte & South, 2026) packages. Base imagery: Esri World Imagery (Esri, 2024). State boundary data: IBGE (Instituto Brasileiro de Geografia e Estatística, 2023).

Sampling employed a repeated-measures design, with control and restoration farms surveyed at baseline (T0) and at successive annual timepoints (T1, T2) following restoration onset (Supplementary Item S1). Sampling effort varied across farms and timepoints (Table 1), generally scaling with farm area (Supplementary Item S2). Several sampling points were removed due to flooding or development and new ones added as restoration progressed.

**Table 1.** Sampling effort by survey method, showing the number of sampling points at each timepoint across each of the eight farms. : baseline (T0), one year after restoration onset (T1), and two years after restoration onset (T2). Point counts were conducted at baseline only. See Supplementary Item S2 for farm area.

| Method | Farm | T0 | T1 | T2 | Total |
| --- | --- | --- | --- | --- | --- |
| Acoustic | Turmalina | 28 | 29 | 29 | 86 |
|  | Tres Irmaos | 14 | 15 | 0 | 29 |
|  | Vale do Candiru | 15 | 16 | 0 | 31 |
|  | Dom Eliseu | 12 | 15 | 0 | 27 |
|  | Uruara | 5 | 13 | 0 | 18 |
|  | Uruara Forest | 3 | 19 | 0 | 22 |
|  | Sao Marcos | 16 | 21 | 0 | 37 |
|  | Nubia | 16 | 0 | 0 | 16 |
|  | **Total** | **109** | **128** | **29** | **266** |
| Camera Trap | Turmalina | 0 | 15 | 14 | 29 |
|  | Tres Irmaos | 9 | 13 | 0 | 22 |
|  | Vale do Candiru | 13 | 13 | 0 | 26 |
|  | Dom Eliseu | 11 | 13 | 0 | 24 |
|  | Uruara | 5 | 11 | 0 | 16 |
|  | Uruara Forest | 3 | 19 | 0 | 22 |
|  | Sao Marcos | 10 | 16 | 0 | 26 |
|  | Nubia | 12 | 0 | 0 | 12 |
|  | **Total** | **63** | **100** | **14** | **177** |
| Point Count | Turmalina | 30 | 0 | 0 | 30 |
|  | Tres Irmaos | 7 | 0 | 0 | 7 |
|  | Vale do Candiru | 0 | 0 | 0 | 0 |
|  | Dom Eliseu | 12 | 0 | 0 | 12 |
|  | Uruara | 11 | 0 | 0 | 11 |
|  | Uruara Forest | 5 | 0 | 0 | 5 |
|  | Sao Marcos | 0 | 0 | 0 | 0 |
|  | Nubia | 0 | 0 | 0 | 0 |
|  | **Total** | **65** | **0** | **0** | **65** |

Pará has a tropical climate: the daily average temperature is 27°C year-round, without much seasonal fluctuation. Data collection occurred between February 2023 and August 2025, spanning both the rainy (December to May, mean rainfall 7.0mm/day) and dry season (June to September, mean rainfall 3.1mm/day) (WorldData.info, 2026).

### Field Data Collection & Processing

To monitor bird activity, three different survey methods were deployed: passive acoustic monitoring, camera traps and point counts. Point counts were conducted at the baseline only, but acoustic monitoring and camera trap surveys were conducted annually. Survey effort varied across methods, with several farms lacking point count or camera trap surveys - an unavoidable caveat due to limited staff resource (Table 1). For acoustic recording, targeting vocalising species, both AudioMoth 1.0.0 and Song Meter Mini recorders were deployed at each sampling point for one week, recording continuously for one minute in every five, operating 24 hours a day (Open Acoustic Devices, 2026; Wildlife Acoustics, 2026; Bradfer-Lawrence et al., 2019). These targeted the frequency band at 48 kHz since this is the frequency most birds vocalise at and were placed two metres above ground. We used BirdNET+ V3.0 for species identification (Kahl et al., 2021; Lasseck et al., 2026).

Camera traps were installed at each sampling point for a seven-day period, following established methods (TEAM, 2011). A mix of camera trap models were used, based on availability: Browning HP5, Bushnell Low Glow, Victure HC300 and Ceyomur CY50. These were placed 10-20 metres from predetermined features (e.g. trails, streams) most likely to capture encounters and mounted 30-50cm above ground. Species were identified through manual review of images, assisted by Wildlife Insights’ AI model (Ahumada et al., 2020; Gadot et al., 2024).

Point counts were conducted by a local ornithologist. These were 10-minute counts of all bird species detected by sight and/or sound within a 200-metre fixed radius and 180-minute window after dawn, as this is the time of maximum bird activity (Bibby et al., 2000; Hutto et al., 1986; dos Anjos et al., 2010; Vielliard et al., 2010). Number of individuals was recorded, with groups considered a single detection. Each sampling point was surveyed four times, across different days.

We compiled and cleaned this data in R (version 4.5.2; R Core Team, 2025) using the *tidyverse* collection of packages (Wickham et al., 2019).

### Acoustic Data Processing

We used BirdNET+ V3.0 developer preview model 3, a pre-release made available in January 2026 (Lasseck et al., 2026), to automatically classify species from the acoustic recordings (all references to “BirdNET” from now onwards refer to BirdNET+ V3.0 unless explicitly stated otherwise). Unlike stable BirdNET releases, this model was subject to change and its performance unestablished, hence the validation procedure below. We ran the standard full-precision (FP32) PyTorch model variant on Imperial’s HPC cluster CX3 (Imperial College Research Computing Service, 2026). We resampled audio files to the model’s native 32 kHz mono format internally and processed them using the default parameters in fixed, non-overlapping 3-second windows. Since BirdNET confidence scores are not directly comparable across species (Wood & Kahl, 2024), no single threshold can be assumed appropriate for all. The standard global *≥*0.5 minimum confidence risks disproportionately excluding rare species given their likely underrepresentation in the training data, which lowers classification reliability (Funosas et al., 2024; Márquez-Rodríguez et al., 2025; Pérez-Granados et al., 2026). We therefore used a lower threshold of 0.1.

### Candidate Species List Generation

To generate a regionally plausible candidate species list, we applied post-processing filtering prior to detection output using a custom checklist of species found within Pará state and a 100 km buffer, based on GBIF, WikiAves and IUCN records (see Supplementary Item S3 for full methods). We then filtered the output detections in R using two separate checklists: the IUCN Red List, restricted to Class Aves and threatened categories (VU/EN/CR) globally, downloaded May 2026 (IUCN, 2026) and the eBird regional list for Pará state (eBird, 2026). Detections were retained only if the species appeared on both lists. We then cross-checked scientific names against the IUCN Red List for taxonomic synonyms or spelling variants using the *stringdist* package (van der Loo, 2014). We merged two duplicate entries, *Porphyrio martinica*/*martinicus* and *Xenops rutilus*/*rutilans*, before combining the lists. We visually inspected eBird distribution maps as a final check to avoid including vagrant records, returning a final list of 23 regionally plausible species for the study site (Table 3). We then validated these candidate species’ detections to confirm their presence.

**Table 3.**
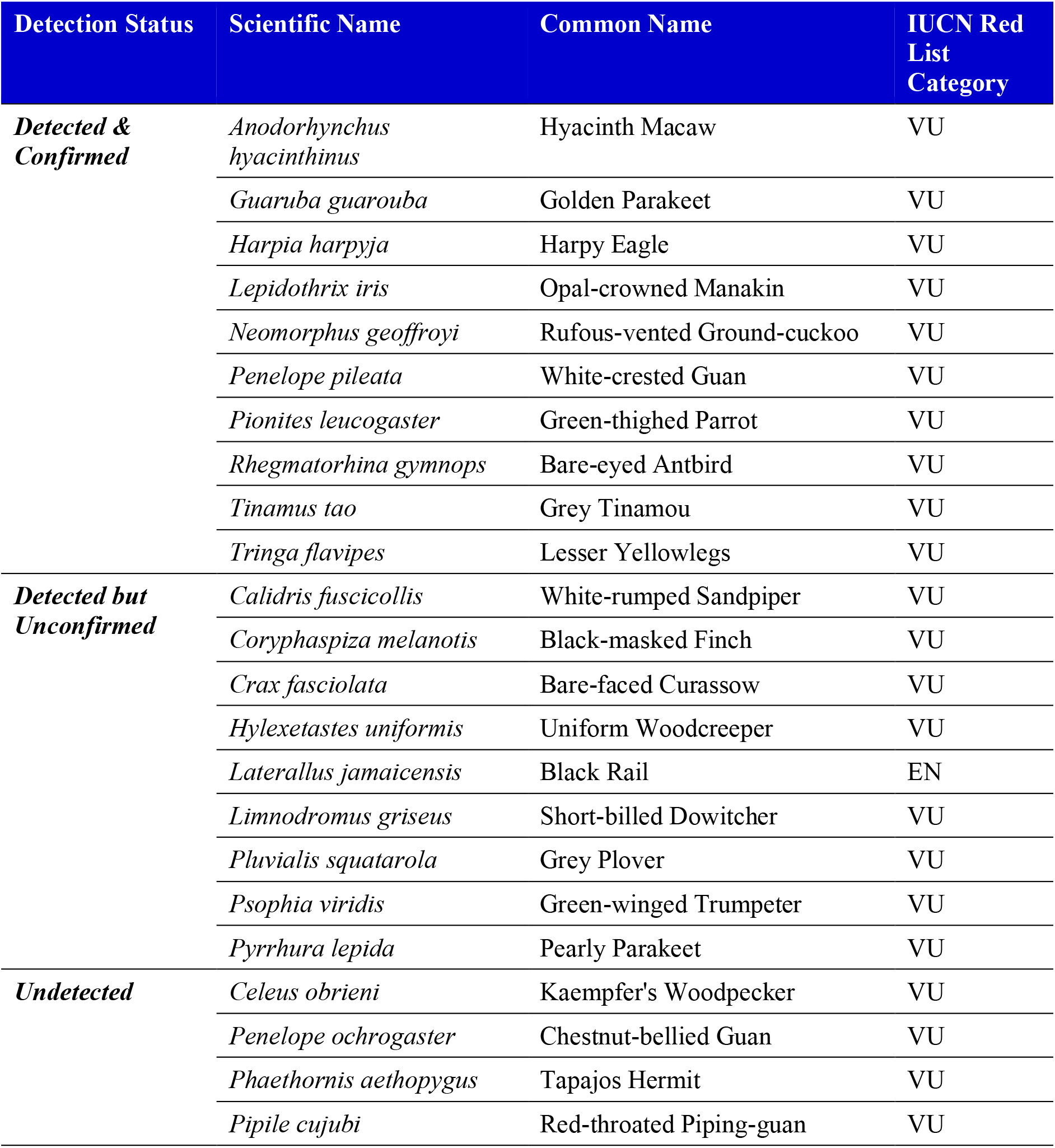
Threatened bird species considered regionally plausible for the study landscape (n = 23), classified under IUCN Red List criteria: Vulnerable (VU), Endangered (EN), Critically Endangered (CR). Regional plausibility determined from two combined sources: a species occurrence list covering confirmed occurrence within Pará state and a 100 km buffer and the eBird regional list for Pará state (see Methods). IUCN status and common names sourced from the global IUCN Red List (IUCN, 2026). Species grouped by detection status: detected and confirmed by validation, detected but unconfirmed (no Tp detections), or undetected by BirdNET.

### Validation Procedure

Manually validating all detections is rarely feasible, so we validated a subset (n=995) by listening to each three-second clip and visually inspecting spectrograms in Audacity (Audacity Team, 2026). The standard approach entails validating detections across confidence score intervals to set species-specific precision thresholds (Tseng, Hodder & Otter, 2025; Wood & Kahl, 2024). This does not transfer well to species with low detection counts (Thompson et al., 2025). Instead, we used an adapted strategy with two different methods, split by species’ detection volume to prioritise validation effort where it was needed most (Chambert et al., 2018). For species with ≥50 detections pre-validation - ‘high-detection’ species, we validated only the single highest-confidence detection at each unique sampling point and timepoint combination (‘occasion’), regardless of the outcome. For species with <50 detections – ‘low-detection’ species, we validated detections exhaustively in descending confidence order until a true positive was found or all detections reviewed. Both approaches assume that higher-confidence predictions are more likely to be correct (Thompson et al., 2025; Wood & Kahl, 2024).

We labelled each detection as a true positive (*Tp*), false positive (*Fp*) or ‘uncertain’ (audible but ambiguous). To inform this, we researched species’ habitat, activity periods, and vocalisations beforehand, including aural training to distinguish candidates from acoustically similar species. We escalated uncertain detections to a local expert ornithologist, whose revalidation was definitive. A subset remained ‘uncertain’ where he was also unable to confirm species identity^1^. These validated detections were then collapsed into a binary presence/absence variable per species, which formed the basis for all subsequent analyses, which we conducted in R.

### Data Analysis

#### BirdNET detection of threatened species

We assessed whether BirdNET can successfully detect threatened species within the study landscape in two stages: firstly, whether BirdNET generated any detections for a candidate species at all; secondly, whether at least one of those detections was a true positive (*Tp*) upon manual validation, confirming the species’ presence at that occasion.

Quantifying model sensitivity/recall, the ability to correctly detect true presences, requires identification of false negatives (Knight et al., 2017; Pérez-Granados, 2023), which is beyond the scope of many studies, this one included. As such, precision is often adopted as a metric of classifier performance or reliability (Bota et al., 2023; Knight et al., 2017). Standard precision is defined as *Tp / (Tp + Fp)*, i.e., true positives divided by total validated detections for a given species. It is typically calculated by validating a subset of detections stratified across confidence bins to ensure representative coverage across the full range (Tseng, Hodder & Otter, 2025; Wood & Kahl, 2024). In this study, we instead calculated precision using only the single highest-confidence detection per occasion (retaining uncertain detections in the total), per the high-detection species’ validation approach. It is therefore likely inflated relative to true precision across the whole confidence range and reflects the reliability of BirdNET’s highest confidence predictions rather than serving as a transferable measure of classifier performance. We therefore term it ‘confirmation rate’ to distinguish from standard precision. We calculated confirmation rate only for high-detection species, since their validation protocol capped *both* true and false detections at one per occasion, producing a consistent, comparable unit. In contrast, the exhaustive validation procedure for low-detection species meant that occasions where false positives outranked the sole true detection in confidence fed cumulatively more false positives into the calculation as a function of search length, systematically lowering confirmation rate for reasons unrelated to classifier reliability.

#### Method comparison: acoustic, camera trap and point count

To assess PAM against traditional methods for detecting threatened species, we compared detections of the 23 regionally plausible candidate bird species across three survey methods: acoustic, camera trap and point count. This was across all eight farms at baseline (T0) only, to allow comparison with point count data. For each species, we aggregated detections by method to farm level, allowing differences in detection performance to be visualised descriptively. We restricted acoustic records to species confirmed by validation, treating camera trap and point count detections as confirmed by default.

#### Testing for temporal change in threatened species presence (T0 – T1)

To test for changes in threatened species’ presence over the first year of restoration (T0 to T1), we compared community-level proportion of sampling points with confirmed presences across the five restoration farms, using a paired t-test to isolate within-farm change over time. We used community-level proportion to account for survey effort differences across farms, calculated for each farm at each timepoint as the total number of confirmed presences across all species retained by validation, divided by the product of sampling points surveyed and number of species (total confirmed/(sampling points surveyed × n species) per timepoint). We ran the test using t.test() (paired=TRUE) and calculated Cohen’s d as the mean paired difference divided by its standard deviation, along with 95% confidence intervals. As a check for temporal effects unrelated to restoration (e.g. climatic variation), we ran an equivalent paired t-test for control farms (n=2; Nubia excluded -T0 data only). Given the limited sample size, this served as descriptive context for the restoration farm result rather than being independently interpretable.

## Results

### BirdNET detection of threatened species

We found that all 23 regionally plausible threatened species were included in the BirdNET label set, indicating that in principle (i.e., if they were present), it was feasible for all 23 to be detected by the classifier within the study landscape. BirdNET detected 82.6% (19/23), 15 of which were high-detection species and 4 low-detection species (Table 3; Figure 2). However, we confirmed only 43.4% (10/23) as truly present: 8 high-detection and 2 low-detection species. We established this by manual validation of 995 total detections across the 19 detected species combined. This means that for 39.1% (9/23) of species, all reviewed detections were false positives. The remaining 17.4% (4/23) of species were not detected by BirdNET at any farm or timepoint.

**Figure 2.**
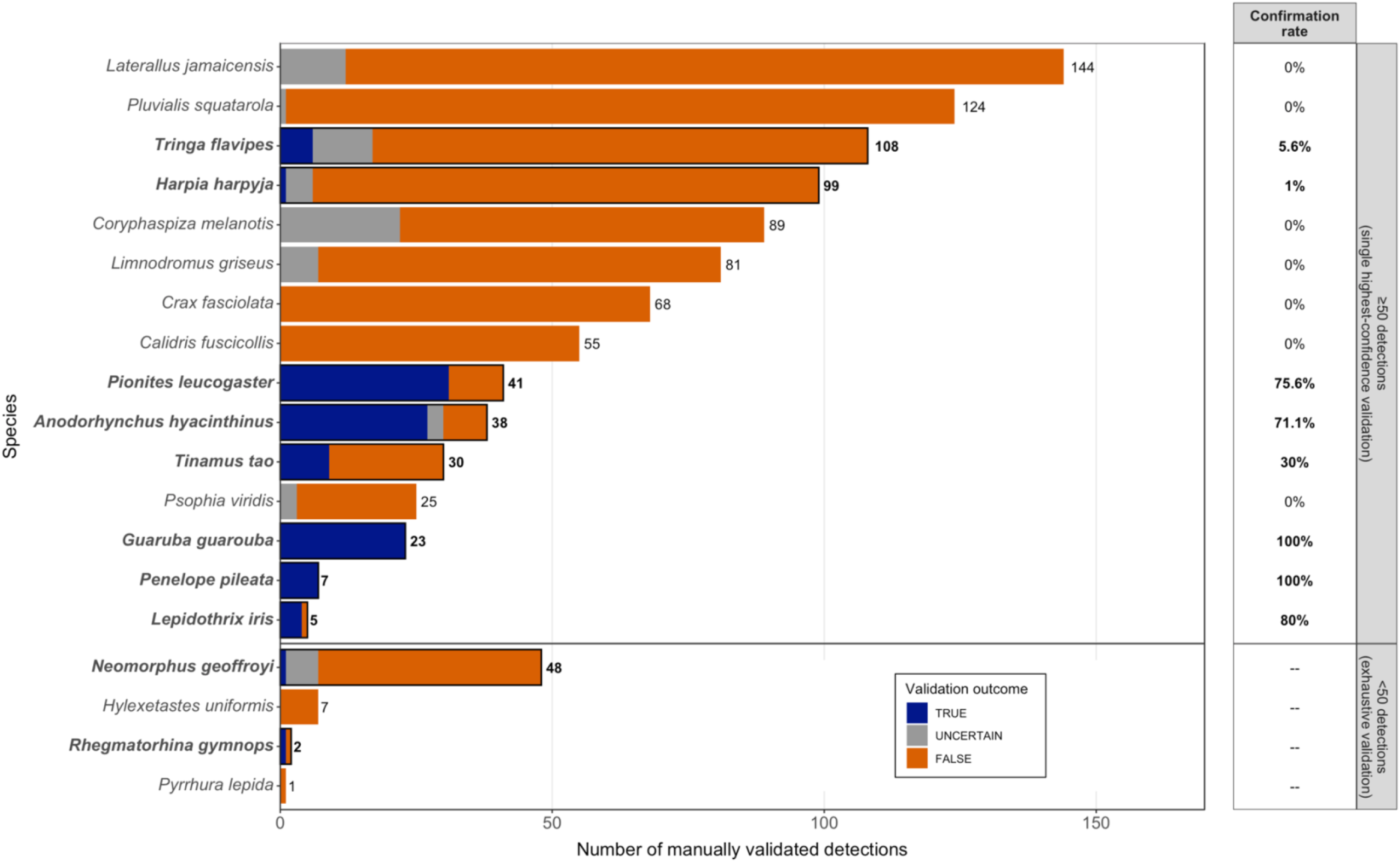
Manual validation outcomes (TRUE/UNCERTAIN/FALSE) and confirmation rates across the 19 detected species (n=995 total reviewed detections). Upper panel: high-detection species (n=15 species) with ≥50 detections pre-validation; lower panel: low-detection species (n=4 species) with <50 total detections pre-validation. Species ordered by validation effort; bar labels show the total number of detections validated. Species with ≥1 confirmed (TRUE) detection are bolded and outlined for clarity. Confirmation rate = % TRUE/Total detections validated for high-detection species; not applicable to low-detection species (denoted ’--’). Each validation attempt is one occasion (sampling point x timepoint) where that species’ single highest-confidence detection was validated. See Supplementary Item S4 for validation outcome breakdown.

Considering only the species BirdNET detected, it confirmed the presence of 52.6% (10/19). Confirmation rate (precision, i.e., the reliability of a species’ detections) could only be calculated for high-detection species under the split validation protocol (see Methods). Among the 15 high-detection species, confirmation rates clustered at both extremes: one third (5/15) at >70%, and the other two thirds (10/15) at ≤ 30%, with none falling in-between. BirdNET detected two species, *Guaruba guarouba* (Golden Parakeet) and *Penelope pileata* (White-crested Guan) with 100% confirmation.

Seven species had confirmation rates of 0%, meaning that all validated detections were false positives, and presence could not be confirmed. This included the two species with the highest number of validated detections – *Laterallus jamaicensis* (Black Rail; n=144) and *Pluvialis squatarola* (Grey Plover; n=124) (see Table 3 for the full list). The remaining three species in this subset were confirmed with confirmation rates of 1-30%. Confirmation rates showed no apparent relationship with validation effort, spanning n=1-144 validated detections across all 19 species. For the four low-detection species for which confirmation rate was not calculable, BirdNET confirmed the presence of 50% (2/4) of these.

### Method comparison: acoustic, camera trap and point count

Across all eight farms at baseline (T0), acoustic monitoring - restricted to confirmed species - detected all species that the traditional methods did, which added no unique species of their own (Figure 3). This constituted 30.4% (7/23) of the threatened species from the regionally plausible list. Camera trap and point count surveys each detected one and two species respectively, with overlap (<10% of plausible species combined). One record of *Penelope pileata* (White-crested Guan) at a single site in one farm (Dom Eliseu) was missed by acoustic monitoring but retrieved by point count. Across all farms pooled, the two species with the highest number of acoustic detections at sampling point level, *Pionites leucogaster* (Green-thighed Parrot; detected at n=15 sampling points) and *Guaruba guarouba* (Golden Parakeet; detected at n=13 sampling points), were completely undetected by camera trap and point count. *Pionites leucogaster* was also the most widely distributed species, detected across n=4 farms.

**Figure 3.**
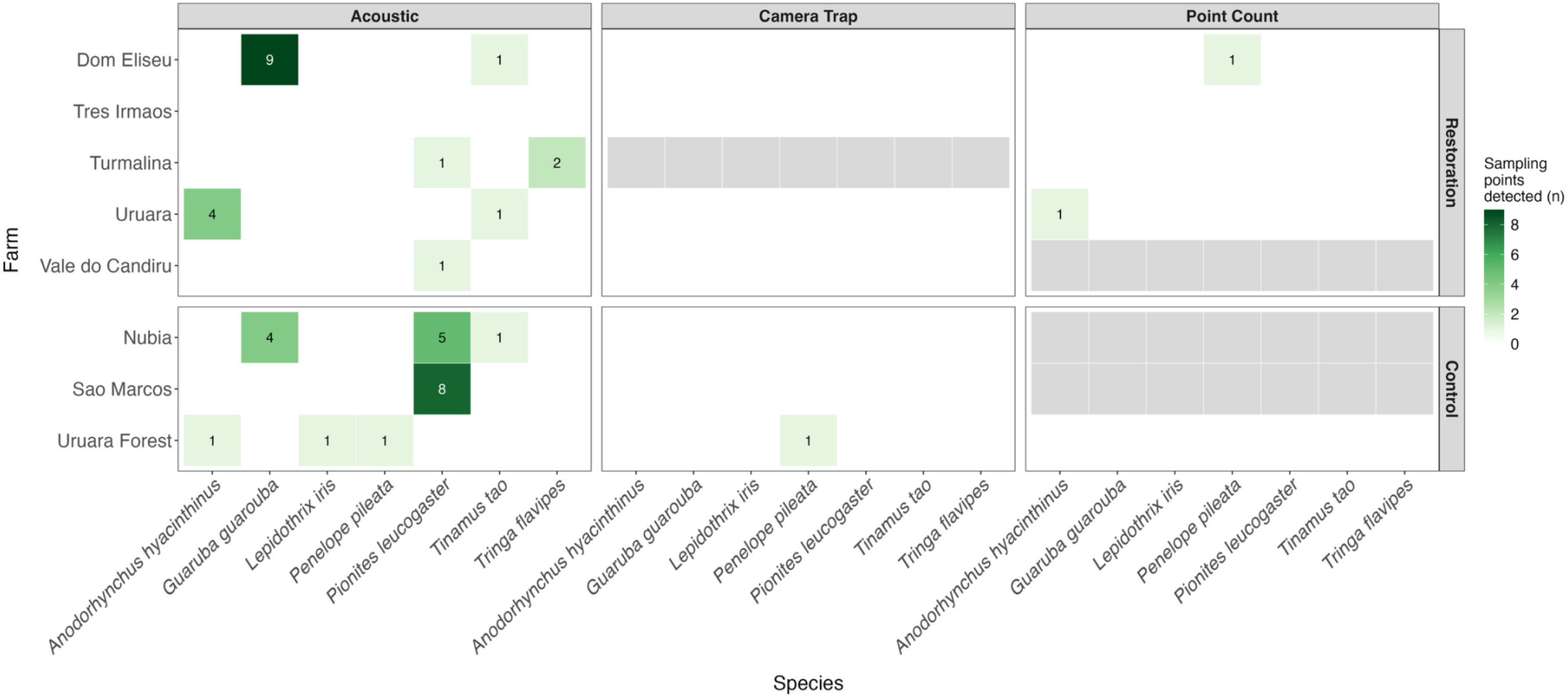
Farms at which each confirmed threatened species was recorded, by survey method (acoustic, camera trap and point count) and farm type (upper panel: restoration; lower panel: control forest). Only the seven species detected by at least one method are displayed. The values displayed in and colour intensity of boxes represent the number of sampling points a species was recorded at within a particular farm (white = 0, dark green = more sampling points). Grey cells with no value displayed indicate farms with no survey effort at baseline (T0), distinct from white cells with no value, which indicate surveyed farms with no records of a given species. Note: baseline survey effort across farms and methods is not uniform (n = 109, 63 and 65 surveyed points for acoustic, camera trap and point count respectively; see Table 1 for full details of sampling effort).

### Testing for temporal change in threatened species presence (T0 – T1)

We found no significant change in community-level presence of threatened species between T0 and T1, a result consistent across both the restoration and control farm t-tests. The mean difference in community-level presence of threatened species between T0 and T1 across restoration farms (n=5 farms) was -0.005 (95% CI [-0.015, 0.006]) (Figure 4A), but the paired t-test showed that this difference was not significant (t(4) = -1.25, p = 0.280, Cohen’s d = -0.56) (Figure 4B). Trajectories were mixed across individual farms, with three farms showing a slight decrease and one a slight increase; one farm remained at zero between both timepoints.

**Figure 4.**
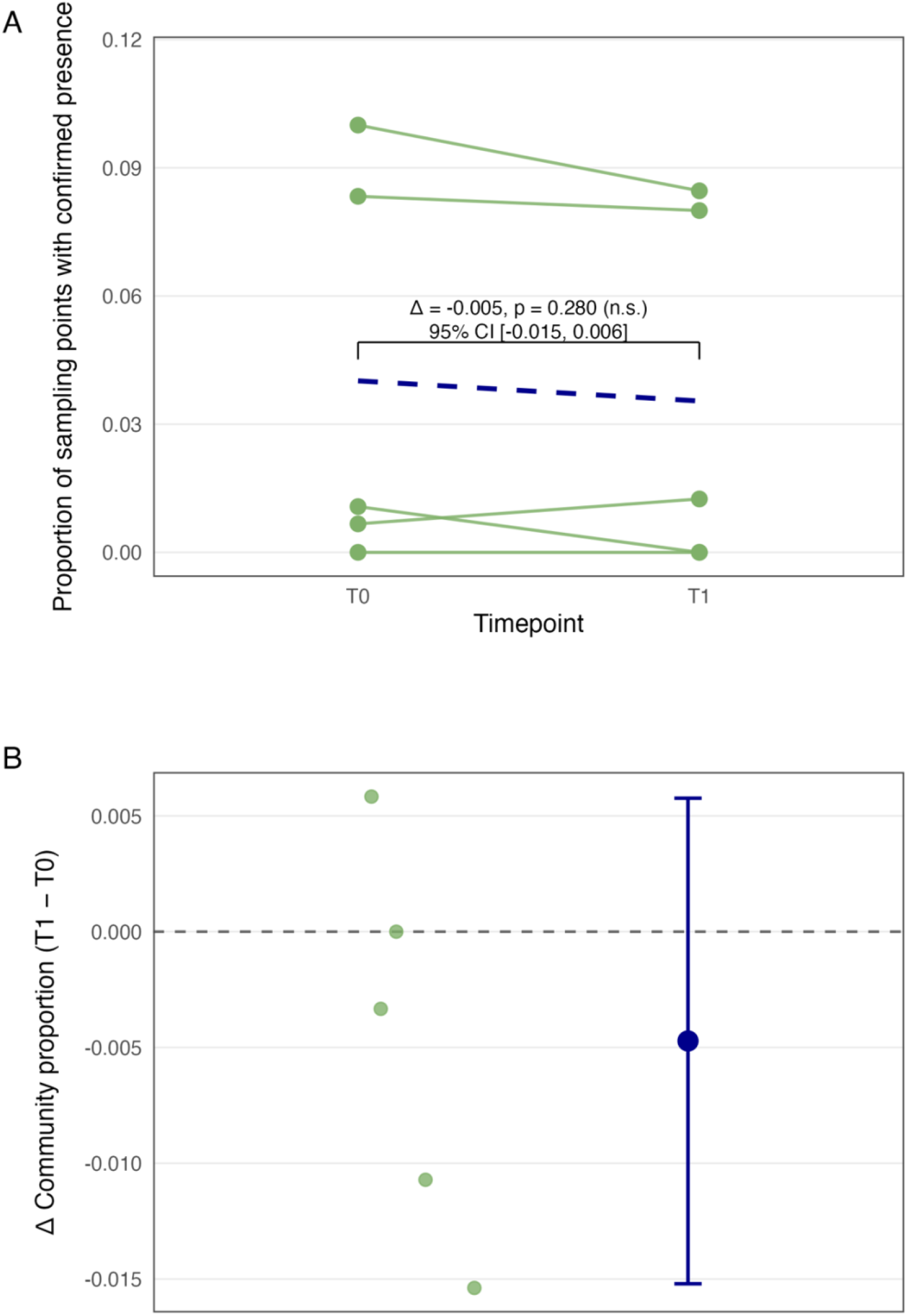
A) Change in community-level proportion of sampling points with confirmed threatened species presence between survey timepoints (T0 = baseline, T1 = first year of active reforestation) for restoration farms. Each green line is an individual farm (n=5). The dashed blue line shows the mean community-level proportion across all five farms at each timepoint: the difference between T0 and T1 is not statistically significant (t(4) = -1.25, p = 0.280). B) Paired change in community-level proportion of confirmed threatened species presence (T1 − T0) per restoration farm (pale green points), with the mean paired difference and 95% confidence interval (in blue). The dashed line marks zero change. The confidence interval spans zero, reflecting the non-significant result.

The control forest paired t-test likewise found no significant difference in community-level presence between T0 and T1 (t(1) = 2.74, p = 0.223), consistent with the restoration result above, although this should be interpreted with caution given its limited power (n=2 farms).

## Discussion

This study set out to establish whether passive acoustic monitoring (PAM) combined with the AI classifier, BirdNET, can support the monitoring of threatened birds in the Brazilian Amazon. To our knowledge, no previous study in this restoration landscape has used PAM with the specific aim of detecting threatened birds, and this is also the first application of BirdNET+ V3.0 in the region. Our results demonstrate that together they can, confirming the presence of ∼43% of the total regionally plausible threatened bird species, including species which had never before been recorded at the study site. PAM markedly outperformed the traditional monitoring methods of camera trap and point counts. However, BirdNET’s classification performance varied across species, and we could not establish whether there were any effects of restoration on threatened species presence within the first year. Nonetheless, this research indicates that PAM combined with BirdNET is a viable and practical method for detecting threatened Amazonian bird species in Brazil.

### BirdNET is effective but performance is species-specific

At the study site, BirdNET confirmed the presence of >40% of the threatened bird species plausible – an encouraging result, given how elusive threatened species are. Whether the remaining species were truly absent from the study site versus present but not confirmed could not be determined from this study.

BirdNET confirmed the presence of just over half of the species it detected, but with mixed performance. For the 15 species for which confirmation rate was calculable, performance grouped at both ends of the scale rather than falling anywhere in-between. Within the context of this restoration site, the model appears capable of reliably detecting a third of species, with confirmation rates above 70%. It performed particularly well for both *Guaruba guarouba* (Golden Parakeet) and *Penelope pileata* (White-crested Guan), which were detected with 100% confirmation. In contrast, it could not reliably detect two thirds of species which had very low confirmation rates, only three of which were confirmed present at the study site at all. The others, with 0% confirmation, were validated without a single true positive.

Such inconsistency in performance is well-documented and affects all classifiers to some degree (Bell et al., 2026; Knight et al., 2017; Wood & Kahl, 2024). Determining the reasons behind misclassification here could help to improve the model’s detection capability (Pérez-Granados, 2023), but in the short-term, high levels of validation are required. This is reflected in the mostly low confirmation rates reported here. While we cannot ascertain whether species with 0% confirmation rates would be confirmed with further validation effort (since 0% may reflect genuine absence), this point is demonstrated by the three confirmed species with confirmation rates below or equal to 30%. For example, *Harpia harpyia* (Harpy Eagle) had a 1% confirmation rate, with only one true positive out of its 99 total detections validated. Since confirming presence is the main goal here, avoiding false negatives matters more than minimising false positives, making this validation burden a necessary cost.

Overall, these results demonstrate that BirdNET has genuine potential for reliably detecting at least a subset of threatened species within Brazilian Amazon restoration sites. However, substantial validation effort is needed if we are to capture *all* occurrences of threatened species possible from the audio.

### PAM detects species missed by traditional monitoring methods

Acoustic monitoring was by far the best method for detecting threatened birds within this restoration site. Out of the total seven threatened species detected across all three monitoring methods combined, acoustic detected all seven, whereas only two of these were detected by camera traps and/or point counts. Additionally, acoustic monitoring provided the first confirmed record at the study site for five species not detected by either traditional method: *Guaruba guarouba* (Golden Parakeet)*, Lepidothrix iris* (Opal-crowned Manakin)*, Pionites leucogaster* (Green-thighed Parrot)*, Tinamus tao* (Grey Tinamou) and *Tringa flavipes* (Lesser Yellowlegs). These species would otherwise have gone unrecorded, with potential downstream conservation implications.

Results of other PAM studies emphasise its success in detecting threatened species. For example, Mosikidi et al. (2023) found 25 previously unrecorded bird species at a site in South Africa, 24 of which were species of IUCN conservation concern. However, they failed to detect several expected and visually confirmed species. This suggests that abandoning traditional monitoring methods altogether would be unwise. Others have found that each method has a distinct taxonomic ‘blind spot’: for example, while camera traps remain valuable for threatened species monitoring more broadly, their strength may lie more in mammals than birds (Doohan et al., 2026; Zwerts et al., 2021) – potentially reflected in this study’s low detection count. Furthermore, that the two species most frequently detected by acoustic monitoring were not detected by camera trap or point count at all suggests that this effect can also operate within taxa, with different detection methods suited to different bird species, depending on their ecology. Point counts also remain useful as an independent dataset for cross-checking PAM results, indicated here by the single *Penelope pileata* (White-crested Guan) detection recovered. A combined approach may therefore be advisable, however, employing multiple methods demands more staff resource, and point counts specifically require identification expertise and local site knowledge (Shonfield & Bayne, 2017).

### No significant effect of restoration in the first year

Contrary to the hypothesised disturbance-driven decrease, we found no significant effect of restoration on community-level threatened species presence between baseline and year one. However, the small sample size (n=5) may have left the t-test underpowered to detect any real effect and may also have masked farm-specific responses to restoration which are lost when pooled at community level. The underpowered control comparison likewise meant that we could not rule out that a broader temporal effect unrelated to restoration explains the null result. Another explanation is that the low confirmed presence at baseline left little scope to register any further decline.

Alternatively, restoration may genuinely not have had an effect, if one year is too short a timeframe for any changes in species presence to emerge. Threatened species, which typically have slow life histories and specific habitat requirements, and thus potentially slower responses to restoration impacts (Uezu & Metzger, 2016), may take even longer to show such changes. As the project is still in its early stages, we should continue to monitor these species and revisit this question as restoration progresses.

### Methodological Limitations

It is important to note that the confirmation rates reported here are mostly derived from low volumes of validated detections (<50 for almost half of species), therefore should be interpreted with caution. While further validation would improve their reliability, they currently offer an initial indication of which species require greater validation effort to confirm presence.

Secondly, survey effort was unequal across monitoring methods: three farms lacked point counts (due to there being only one on-site expert), and one farm lacked camera trap surveys (Table 1). This may partly explain the higher number of species detected by acoustic monitoring, as it is unclear whether the other methods would have detected the same species at these farms. Future comparisons should ensure that survey effort is standardised, for example, during field protocol design. Nevertheless, the imbalance here highlights the value of PAM as a low-effort survey method that enabled monitoring which otherwise would not have happened.

### Conservation Implications

PAM and BirdNET can meaningfully contribute to threatened species conservation. Together, they have reinforced the conservation value of this restoration landscape by confirming the presence of ten threatened species across these farms. Four of these species are Brazilian endemics, and *Guaruba guarouba*, the Golden Parakeet, is contender for Brazil’s national bird, making it a flagship species as well as an Important Bird Area (IBA) trigger species (de Moraes et al., 2025; Pacheco et al., 2021; Birdlife International, 2018; Laranjeiras, 2020). If further years of monitoring confirm persistence of these species and can provide information on their population sizes, this could help secure further funding for restoration projects like this one.

Furthermore, these findings carry policy and management implications. The five species that were undetected by both traditional monitoring methods represent the first confirmed record of their presence at the study site. Without PAM, these would have falsely been treated as absent – a result which would have informed future restoration development plans and ecological research, with potentially drastic consequences for their populations at these farms.

### Conclusions & Future Direction

PAM, combined with AI classification by BirdNET+ V3.0, shows real potential for threatened species monitoring in Brazilian Amazon restoration. This potential is strongest for two species with 100% confirmation rates; however, the majority require manual validation.

This raises a practical question: is using BirdNET to detect threatened species worth it, given the high validation burden? This study reviewed 995 detections, output by BirdNET as 3-second clips. This is equivalent to a minimum of ∼50 minutes of validation, although in practice this took several hours, since calls often cut across multiple clips and the validators had to listen more than once to identify them. The alternative would have been to conduct PAM without any AI classifier, instead relying on trained ornithologists to identify species from recordings (Cole et al., 2022; Winiarska, Szymański & Osiejuk, 2025). Based on the recording schedule (1 minute in every 5, 24 hrs/day, for 7 days), this would have required listening to approximately 8,940 hours of audio across the eight study farms, equivalent to an estimated 1,117 8-hour working days, which is impractical. Furthermore, the primary validator conducted this validation from England, having never visited Brazil or South America. This demonstrates that non-specialists, with no prior familiarity with the study region or its species, can successfully learn to identify species from recordings and confirm their presence at a site retrospectively - highlighting the feasibility of using AI classifiers like BirdNET for these purposes. While still labour-intensive, this effort is necessary if threatened species are to be detected at all.

Compounding this, our results here have shown that detecting rare species requires a different validation approach from the convention for common species, given their low detection counts. Ideally, every detection would be validated to ensure that all true occurrences present in audio recordings are captured, producing the most complete and accurate record of species presence. However, for situations where time and resource are limited, here we have demonstrated a legitimate alternative: validating the single highest-confidence detection per occasion. By concentrating on the predictions which are most likely to be correct (Thompson et al., 2025), this approach prioritises the question that matters most to conservation practitioners: whether or not a species is present.

Furthermore, unlike confidence thresholds, which are context-specific (Wood & Kahl, 2024), confirmed presences are a permanent, reusable record. This validation method is transferable to other AI classifiers (e.g., BatDetect2 (Aodha et al., 2022), Perch 2.0 (Merriënboer et al., 2026), custom CNNs) and across any taxonomic group. Beyond threatened species detection, we also recommend it as a practical alternative for any species that do not meet the ∼50 detections that conventional confidence-stratified validation approaches require.

In conclusion, PAM with classification by BirdNET, in combination with the validation method developed here, offers a practical and cost-effective approach for detecting threatened species across tropical forest restoration landscapes. Although validation currently remains a necessity for threatened species, model advances should reduce the need for this and eventually lead to a fully automated monitoring approach. Governments and conservation funding bodies should therefore invest more in developing BirdNET and other emerging AI classifiers to meet the standards required for robust monitoring of such species. This investment is especially needed in the tropics, where performance of these models lags behind that achieved in the Global North (Sethi et al., 2024). Research diagnosing the main drivers of misclassification within the threatened species community, and finding practical solutions to these, would also be useful. For example, previous studies have raised concerns about a shortage of training data (Bell et al., 2026; Sethi et al., 2024), which is a direct consequence of these species’ rarity and therefore may not be resolvable by simply collecting more data. Such progress would benefit a wide range of stakeholders, from governments and conservation bodies to carbon credit organisations. Ultimately, it could extend AI-assisted PAM beyond validation-reliant presence detection to lower-effort, reliable monitoring of threatened species at scale – tracking the impacts of forest restoration on these species as it progresses.

## Supporting information

Supplementary Item S1

Supplementary Item S2

Supplementary Item S3

Supplementary Item S4

## Acknowledgements

The study site is managed by the Brazilian carbon removal company, Mombak. Acoustic and camera trap data collection in the field was conducted by Dr Jenna Lawson, Raimundo Ferreira, Sebastian Pipins and PhD students at the Imperial College London Community and Landscape Ecology (CaLE) Lab: Rhys Preston-Allen and Xiuhan Zhang. Point count data was collected by Dr Celso Parruco. Rhys Preston-Allen was responsible for species identification from camera trap data. Dr Celso Parruco assisted with species identification from acoustic data by providing a secondary validation of extracted BirdNET clips.

Barbara Lima Silva (PhD, CaLE Lab) was responsible for compilation of the Pará + 100 km buffer species list.

Computational resources, specifically use of the CX3 High Performance Computing (HPC) facility, were provided by the Imperial College Research Computing Service (http://doi.org/10.14469/hpc/2232). Rhys Preston-Allen assisted with the task of running the HPC to extract BirdNET detections.

## Data and Code Availability

Data and code are available at this repository, currently set to private to protect sensitive location information on IUCN Red List threatened species. Access available upon request.

https://github.com/heatherfortune/FortuneHX_MRes_Thesis_2026

Raw audio data is sensitive and therefore available upon request from Professor Cristina Banks-Leite.

## Footnotes

1 At the time of submission, a subset of detections remains pending secondary validation. Results should be interpreted with this in mind.

