## Supplementary Item S1 for "Towards Automated Acoustic Monitoring of Threatened Birds in Tropical Forest Ecosystem Restoration"

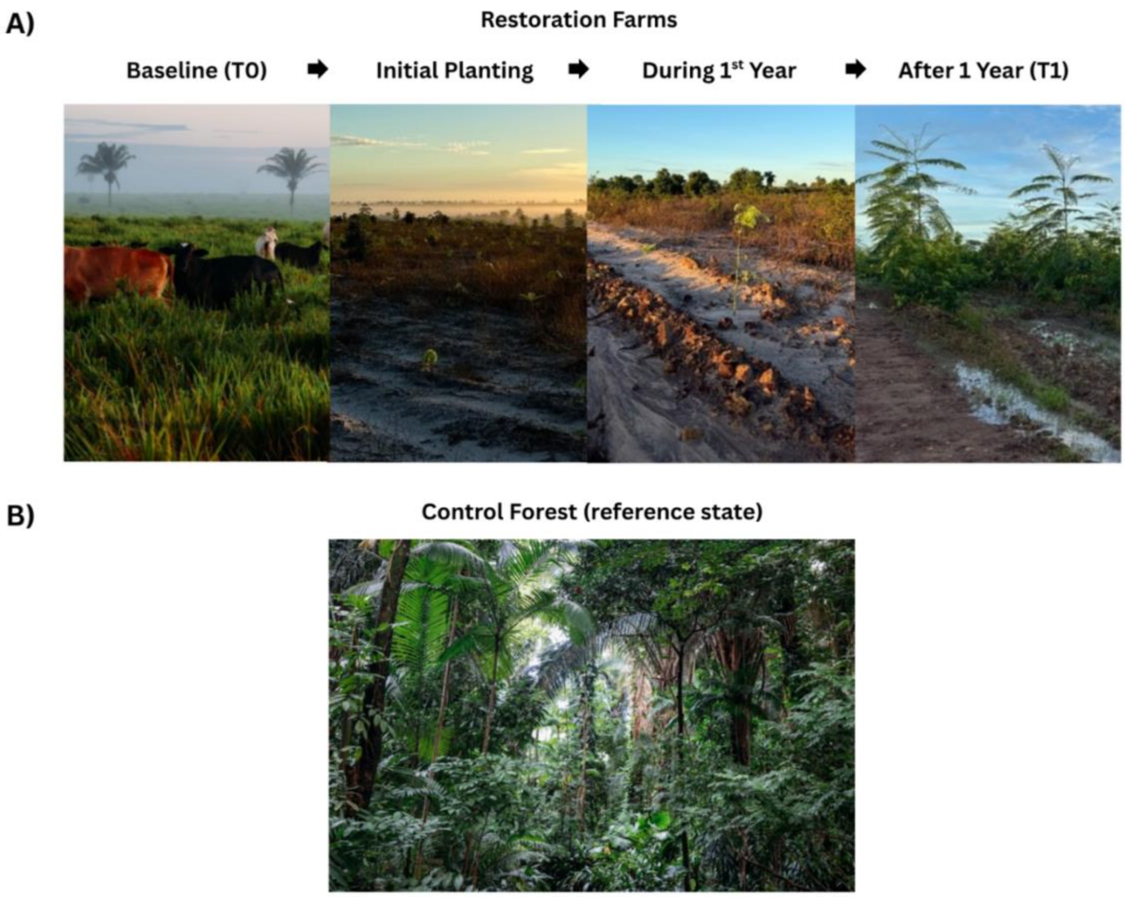


***Figure S1.*** *Photographs from the Pará study site showing general conditions of farms at different stages of restoration (A), compared to a control forest site (B). Photo credits: Rhys Preston-Allen & Xiuhan Zhang.*
