## Supplementary Item S2 for "Towards Automated Acoustic Monitoring of Threatened Birds in Tropical Forest Ecosystem Restoration"

***Table S2.*** *Approximate area (ha) of study restoration and control farms.*

| | Farm | Total Area (ha) |
|---|---|---|
| ***Restoration*** | Turmalina | 2906 |
| | Tres Irmaos | 669 |
| | Vale do Candiru | 1482 |
| | Dom Eliseu | 4000 |
| | Uruara | 3000 |
| ***Control*** | Uruara Forest | 7000 |
| | Sao Marcos | 3760 |
| | Nubia | 4000 |
