## Supplementary Item S3 for "Towards Automated Acoustic Monitoring of Threatened Birds in Tropical Forest Ecosystem Restoration"

***Item S3.*** *Pará + 100km buffer species list extraction methods, written by Barbara Lima Silva. Script also created and run by Barbara Lima Silva.*

This details the methods used to process species occurrence data from three different sources: GBIF, WikiAves, and IUCN. It aims to standardize the data, create a comprehensive comparison of species, analyze their occurrences, and visualize the results through various plots.

**File Paths**

- Defined file paths for the CSV files containing species data from different databases:
  - GBIF: Unique species occurrence data.
  - WikiAves: Bird species data.
  - IUCN: Species range data from the International Union for Conservation of Nature.

**Reading CSV Files**

- Loaded the species data from the defined file paths into three separate DataFrames:
- gbif: Holds data from GBIF.
- wiki: Holds data from WikiAves.
- iucn: Holds data from IUCN.

**Standardizing Columns**

- Selected relevant columns for analysis: phylum, class, family, genus, species, status, reference.
- Standardized text formatting:
- Stripped whitespace from species and genus names.
- Capitalized family names and class names for consistency.

**Merging Species Data**

- Combined data from GBIF, WikiAves, and IUCN into a single DataFrame called all_species.
- Removed duplicate species entries based on the species name.
- Records not identified to species level (e.g., 'Genus sp.') were removed prior to analysis.
- Created a taxonomy DataFrame containing only the taxonomic classification of the combined unique species.

**Creating Presence Matrix**

- Created a comparison DataFrame to show the presence of species across the three databases:
- Added binary indicators (X or empty) for each source to indicate whether a species is present in GBIF, WikiAves, or IUCN.

**Calculating Percentage by Group**

- Calculated the total number of unique species and grouped them by class.
- Computed the percentage contribution of each vertebrate group relative to the total species count.

**Visualization - Bar Chart**

- Created a bar chart to visualize the percentage of total species contributions from each vertebrate group.
- Annotated the bars with the number of species represented.

**Visualization - Pie Chart**

- Generated a pie chart to illustrate the relative composition of vertebrate groups across all databases combined.

**Saving Comparison Data**

- Exported the comparison DataFrame to a CSV file for future reference.

**Total Species Count**

- Calculated and printed the total number of species for each database:
    - GBIF
    - WikiAves
    - IUCN

**Intersection of Species Lists**

- Identified and printed the number of species present in multiple databases:
- Intersection between GBIF and IUCN.
- Intersection between GBIF and WikiAves.
- Intersection between IUCN and WikiAves.
- Species present in all three databases.

**Visualization - Total Species Comparison**

- Created a bar chart to compare the total species richness across the three databases.

**Species Count by Class**

- Grouped species by class for each database and created a summary DataFrame to show species counts by class.

**Visualization - Species by Class**

- Generated a bar chart to visualize species richness by class across the three data sources.

**Saving Class Data**

- Exported the class summary DataFrame to a CSV file.

**Checking for Duplicates**

- Checked and printed the number of duplicate species entries for each database.
- Identified and printed the unique duplicated species from GBIF.

**Status Comparison by Source**

- Created a status table to compare the conservation status of each species across the three databases.
- Mapped the status of each species using the status columns from each source.

**Detecting Status Divergence**

- Implemented a function to check for divergence in species status across the databases.

- Identified and counted species with differing status across the three sources.

**Saving Status Comparison Data**

- Exported the status comparison DataFrame to a CSV file for further analysis.

**Preparing Data for Status Visualization**

- Created a function to count the number of extant and possible extant species by class from each database.

**Visualization - Status by Group**

- Generated a grouped bar chart to visualize the occurrence status of species by vertebrate group.

**Divergence Analysis**

- Analyzed and printed the number of species with status divergence by class.
- Calculated the total number of species per class and the percentage of divergent species.

**Saving Divergence Summary**

- Created a summary DataFrame to store total species, divergent species, and percentage divergence by class.
- Exported the divergence summary to a CSV file for documentation purposes.

**Key Technical Specifications:**

- Data Sources:
  - GBIF: Global Biodiversity Information Facility
  - WikiAves: Brazilian bird observation platform
  - IUCN: International Union for Conservation of Nature
- Data Operations:
  - Data cleaning and standardization.
  - Merging and deduplication of species records.
  - Creation of presence/absence matrices.
  - Visualization through bar and pie charts for comparative analysis.
- Analysis:
  - Calculation of species richness and comparison across databases.
  - Analysis of species status (extant and possible extant) and detection of discrepancies.
