## Supplementary Item S4 for "Towards Automated Acoustic Monitoring of Threatened Birds in Tropical Forest Ecosystem Restoration"

***Table S4. Manual validation outcomes across the 19 species detected by BirdNET, by raw count for each validation category (TRUE, FALSE and UNCERTAIN).*** *High-detection species (≥50 total detections pre-validation) had only the single highest-confidence detection validated per occasion; low-detection species (<50 total detections pre-validation) were validated exhaustively in order of descending confidence until a true positive was found or all candidate detections had been reviewed (see Methods). Species with ≥1 confirmed (TRUE) detection are bolded for clarity. Species ordered by validation effort.*

| Species | Detection Class | TRUE | FALSE | UNCERTAIN | Total Reviewed |
|---|---|---|---|---|---|
| *Laterallus jamaicensis* | high | 0 | 132 | 12 | 144 |
| *Pluvialis squatarola* | high | 0 | 123 | 1 | 124 |
| ***Tringa flavipes*** | **high** | **6** | **91** | **11** | **108** |
| ***Harpia harpyja*** | **high** | **1** | **93** | **5** | **99** |
| *Coryphaspiza melanotis* | high | 0 | 67 | 22 | 89 |
| *Limnodromus griseus* | high | 0 | 74 | 7 | 81 |
| *Crax fasciolata* | high | 0 | 68 | 0 | 68 |
| *Calidris fuscicollis* | high | 0 | 55 | 0 | 55 |
| ***Pionites leucogaster*** | **high** | **31** | **10** | **0** | **41** |
| ***Anodorhynchus hyacinthinus*** | **high** | **27** | **8** | **3** | **38** |
| ***Tinamus tao*** | **high** | **9** | **21** | **0** | **30** |
| *Psophia viridis* | high | 0 | 22 | 3 | 25 |
| ***Guaruba guarouba*** | **high** | **23** | **0** | **0** | **23** |
| ***Penelope pileata*** | **high** | **7** | **0** | **0** | **7** |
| ***Lepidothrix iris*** | **high** | **4** | **1** | **0** | **5** |
| ***Neomorphus geoffroyi*** | **low** | **1** | **41** | **6** | **48** |
| *Hylexetastes uniformis* | low | 0 | 7 | 0 | 7 |
| ***Rhegmatorhina gymnops*** | **low** | **1** | **1** | **0** | **2** |
| *Pyrrhura lepida* | low | 0 | 1 | 0 | 1 |
